# Leveraging engineered *Pseudomonas putida* minicells for bioconversion of organic acids into short-chain methyl ketones

**DOI:** 10.1101/2024.01.06.574483

**Authors:** Ekaterina Kozaeva, Manuel Nieto-Domínguez, Kent Kang Yong Tang, Pablo Iván Nikel

## Abstract

Methyl ketones, key building-blocks widely used in diverse industrial applications, largely depend on oil-derived chemical methods for their production. Here, we investigated bio-based production alternatives for short-chain ketones, adapting the solvent-tolerant soil bacterium *Pseudomonas putida* as a host for ketone biosynthesis either by whole-cell biocatalysis or using engineered minicells, chromosome-free bacterial vesicles. Organic acids (acetate, propanoate and butyrate) were selected as the main carbon substrate to drive the biosynthesis of acetone, 2-butanone and 2-pentanone. Pathway optimization identified efficient enzyme variants from *Clostridium acetobutylicum* and *Escherichia coli*, which were tested under both constitutive and inducible expression of the cognate genes. By implementing these optimized pathways in *P*. *putida* minicells, which can be prepared through a simple 3-step purification protocol, the feedstock was converted into the target short-chain methyl ketones, remaining catalytically functional for >4 months. These results highlight the value of combining morphology and pathway engineering of non-canonical bacterial hosts to establish alternative bioprocesses for toxic chemicals that are difficult to produce by conventional approaches.

**GRAPHICAL ABSTRACT:** 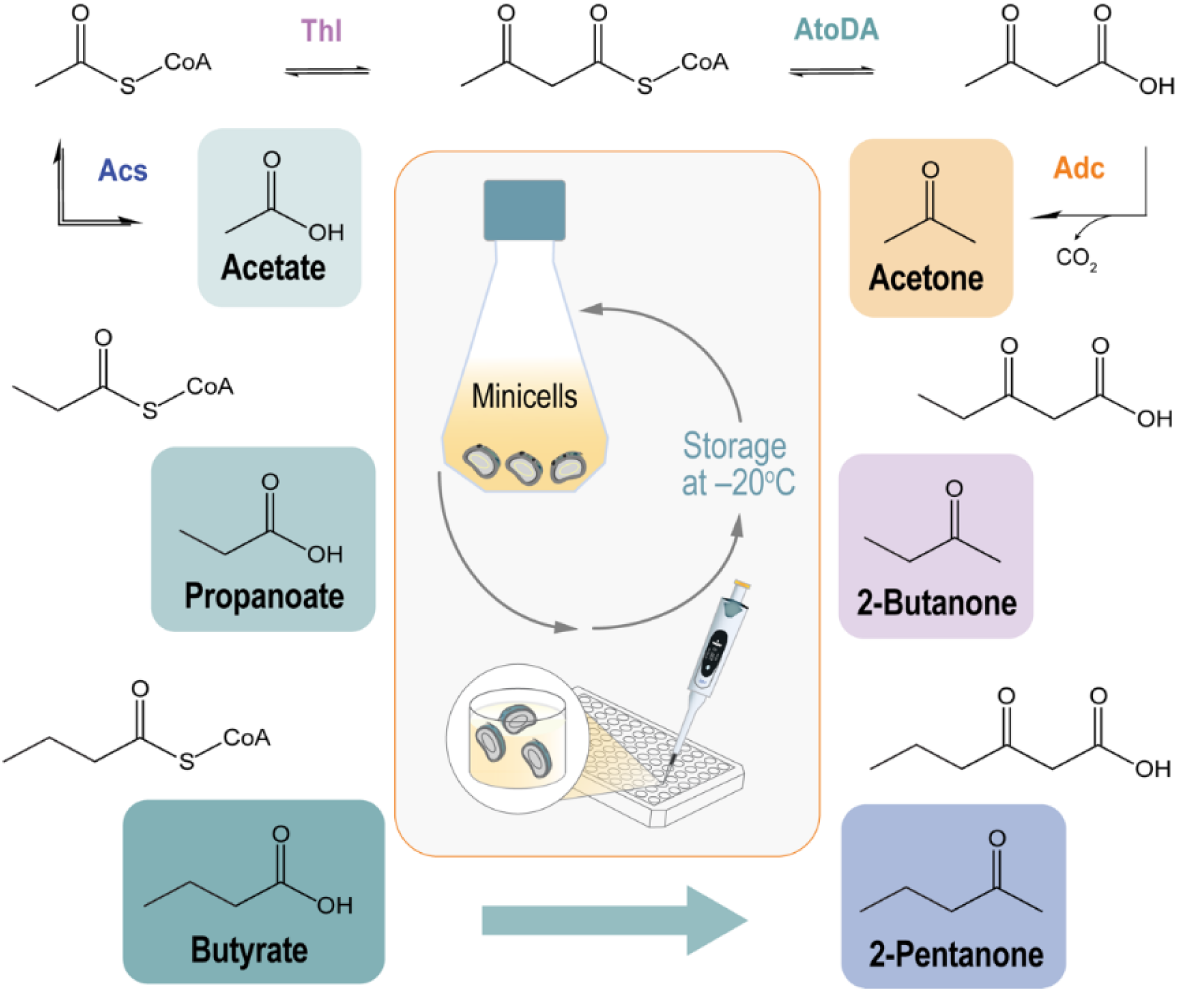

## INTRODUCTION

Ketones, a class of structurally-diverse molecules with the general formula *R*—C(=O)—*R*’, where *R* and *R*’ can be a variety of carbon-containing substituents, are currently produced using traditional, oil-based chemistry [1]. Methyl ketones (MKs), a subset of this family of compounds where one ligand on the carbonyl carbon is a CH_3_ group, play an important commercial role. Because of their strong solvent properties and generally high evaporation rates, ketones are used as building-blocks in the fragrance, flavor, textile, pesticide and agrochemical industries [2,3], as well as being key precursors for the synthesis of pharma molecules [4]. 2-Pentanone [CH_3_—(CH_2_)_2_—C(=O)—CH_3_], a prime example of the broad MKs family, has multiple applications in the fragrance sector and it has been recognized as a potent inhibitor of prostaglandin production associated with colon carcinogenesis [5]. Hence, greener alternatives for ketone production are needed to fulfil the growing market demands, and the adoption of robust microbial cell factories emerge as an attractive option to this end [6,7]. Toxicity issues, imposed on the producer cells by either intermediates or products of solvent biosynthesis pathways, continue to be a major hurdle that impairs the development of robust bioprocesses [8,9]. This occurrence frequently leads to the loss of plasmid(s) encoding components of the biosynthesis pathways or the accumulation of mutations in the genes thereof, resulting in non-producing phenotypes [10–12].

Several strategies have been implemented towards extending the lifespan of actively producing cell factories, yet growth-coupled bioprocesses involving harsh, reactive intermediates and products tend to display limited yields due to the stability issues listed above. An alternative approach that can help overcoming such limitations is using anucleated, bacteria-derived vesicles as biocatalysts [13,14]. Chromosome-free minicells, for instance, naturally occur within bacterial populations albeit at very low frequencies [15]. Under normal growth conditions, the *Z* ring constricts and recruits the peptidoglycan synthesis machinery together with its associated proteins [16]. The dynamic assembly of FtsZ units in rdo-shaped bacteria is controlled by the Min system, which encompasses the MinC, MinD and MinE structural proteins [17]. MinC and MinD inhibit FtsZ assembly, whereas MinE acts as a negative regulator of MinCD. Minicells are formed when there is a limited abundance of MinC and MinD, whereupon FtsZ buildup promotes asymmetric cell division [18]. Unlike their parental counterparts, minicells do not contain nucleic acids and are unable to further divide; for this reason, they have historically been exploited as a model for studying protein synthesis [19]. In spite of their recognized potential as (mini)cell factory [20], these chromosome-free bacterial vesicles have not been extensively exploited for bioproduction. Enhanced protein production, for instance, has been demonstrated in *Escherichia coli* minicells [21]—since plasmids are preferentially located at the poles of the parental bacterium [22], their enrichment in minicells is facilitated *via* either active partitioning or random distribution. Moreover, resources in the minicells (e.g. reducing power and energy equivalents) are expected to be allocated to bioproduction instead of housekeeping processes, e.g. genome maintenance and duplication. Among the microbial *chassis* implemented for metabolic engineering thus far, minicell applications have been exploited mostly for *E. coli* and *Bacillus subtilis* [19]. No attempts have been reported on producing these anucleated vesicles from *Pseudomonas putida*, a non-pathogenic soil bacterium extensively adopted for microbial engineering applications [23–26].

Building on the well-known solvent tolerance features of *P. putida* [27–29] and its ability of using a wide range of carbon sources, we have engineered genome-reduced *P. putida* strains for biosynthesis of short-chain (C_3_-C_5_) (methyl) ketones from organic acids, with a focus on 2-pentanone. The capacity of *Pseudomonas* species to produce long-chain MKs is illustrated in a superb example of model-guided engineering of *P*. *taiwanensis* VLB120 [30], which enabled the synthesis of C_11_-C_17_ MKs using sugars as the substrate. In our current study, the tolerance of *E. coli* and *P. putida* to products and substrates was compared, and production conditions were optimized for whole-cell 2-pentanone biosynthesis by using butyrate as the main carbon source. Different synthetic pathway designs were likewise tested, analyzing the performance of constitutive and inducible expression systems across production conditions. Furthermore, we explored the potential of *P*. *putida* minicells for chemical production, adopting MKs as model compounds known to exert toxicity in bacterial cells. Thus, we engineered *P. putida*-derived minicells to express selected pathway variants, demonstrating production of acetone, 2- butanone and 2-pentanone from acetate, propanoate and butyrate, respectively. Programmable *P*. *putida* minicells had the ability to stably produce ketones over extended timeframes (up to 4 months)— a first case example of an ‘off-the-shelf’ cell factory for MK biosynthesis. This strategy underscores the potential of integrating synthetic morphology with pathway engineering, offering innovative approaches for sustainable ketone biosynthesis.

## RESULTS AND DISCUSSION

### Exploring the potential of P. putida for short-chain ketone biosynthesis from organic acids

A major challenge for efficient short-chain ketones production in engineered bacterial cell factories is the stress caused by the endogenously produced chemicals, which (similarly to other solvents) [31] could inhibit growth or even cause cell death [32]. While acetone (2-propanone, C_3_) and 2-butanone (C_4_) biosynthesis by engineered microorganisms has been explored in the primary literature [33–36], reports on bio-based approaches for 2-pentanone (C_5_) production are scarce. A study by Lan et al. [37] highlighted product toxicity as the key factor severely impairing 2-pentanone biosynthesis by engineered *E. coli* strains using glucose as a carbon source. The growth of *E. coli* JCL299, one of such modified strains [37], was reported to be inhibited by 50% in the presence of as little as 0.6 g L^−1^ 2- pentanone, and bacterial growth was fully arrested when the concentration of the ketone increased to 5 g L^−1^. As *P. putida* is known to be a solvent-tolerant host, we explored its capacity for short-chain MK production, with 2-pentanone as a proxy of this family of compounds.

As a first step in testing the performance of *P. putida* as a production platform, we examined the effect of adding 2-pentanone to cultures of *P. putida* SEM1.3. Strain SEM1.3 (**Table 1**) is a refactored derivative of *P. putida* EM42 [38] a reduced-genome version of the platform strain KT2440 [39]. The modifications introduced in *P. putida* SEM1.3 comprise (i) deletion of the *benABCD* gene cluster [40] to abolish oxidation of 3-methylbenzoate (3-*m*Bz), allowing the use of this molecule as a gratuitous (i.e. non-metabolizable) inducer of the XylS/*Pm* expression system without the interference to optical density at 600 nm (OD_600_) measurements typically caused by brown-colored catechols and other products of aromatic compound metabolism [41] and (ii) elimination of the native *pha* gene cluster, *phaC1ZC2DFI* [42] to avoid any potential metabolic cross-talk that could compete for acetyl-coenzyme A (CoA), key precursor for short-chain ketone biosynthesis *via* acetyl-CoA–dependent chain elongation. *P. putida* SEM1.3 was grown either in rich lysogeny broth (LB) or de Bont minimal (DBM) medium supplemented with 1% (w/v) glucose as the main carbon source and spiked with 2-pentanone at different concentrations (i.e. 1, 2.5, 5, 10, 25 and 50 g L^−1^). Under all conditions tested, the impact of 2- pentanone on cell physiology was more evident when cells grew in a mineral medium than in a rich broth. Bacterial growth, assessed as the final OD_600_ values after 24 h, started to be affected at ketone concentrations above 5 g L^−1^, but bacterial growth was still observed with up to 25 or 50 g L^−1^ of 2- pentanone both in DBM medium or LB medium, respectively (**Fig. 1A**). As an example, the final cell density was only reduced to ca. 40% in LB medium spiked with 2-pentanone at 50 g L^−1^. In general, *P. putida* SEM1.3 fared significantly better under all of these conditions when compared with data reported for *E. coli* JCL299 [37]. These results illustrate the ability of *P. putida* to adapt to high solvent concentrations, a phenotype that could be mediated by modifications in the surface of the outer membrane and other natural stress response mechanisms [27]. Furthermore, we explored whether *P. putida* can utilize 2-pentanone as a carbon source by incubating strain SEM1.3 in DBM medium supplemented with 1 g L^−1^ of 2-pentanone as the sole carbon source (i.e. a concentration that should not mediate any toxicity) and assessing bacterial growth over 100 h (**Fig. 1B**). Control cultures with 1 g L^−1^ glucose were run in parallel to benchmark growth patterns, reaching full saturation within 24 h of incubation (**Fig. 1B**). No growth was detected when strain SEM1.3 was incubated with the ketone, indicating that *P. putida* is unable to utilize 2-pentanone as a carbon substrate, underscoring the potential of this bacterial platform for short-chain MK bioproduction.

**Fig. 1.**
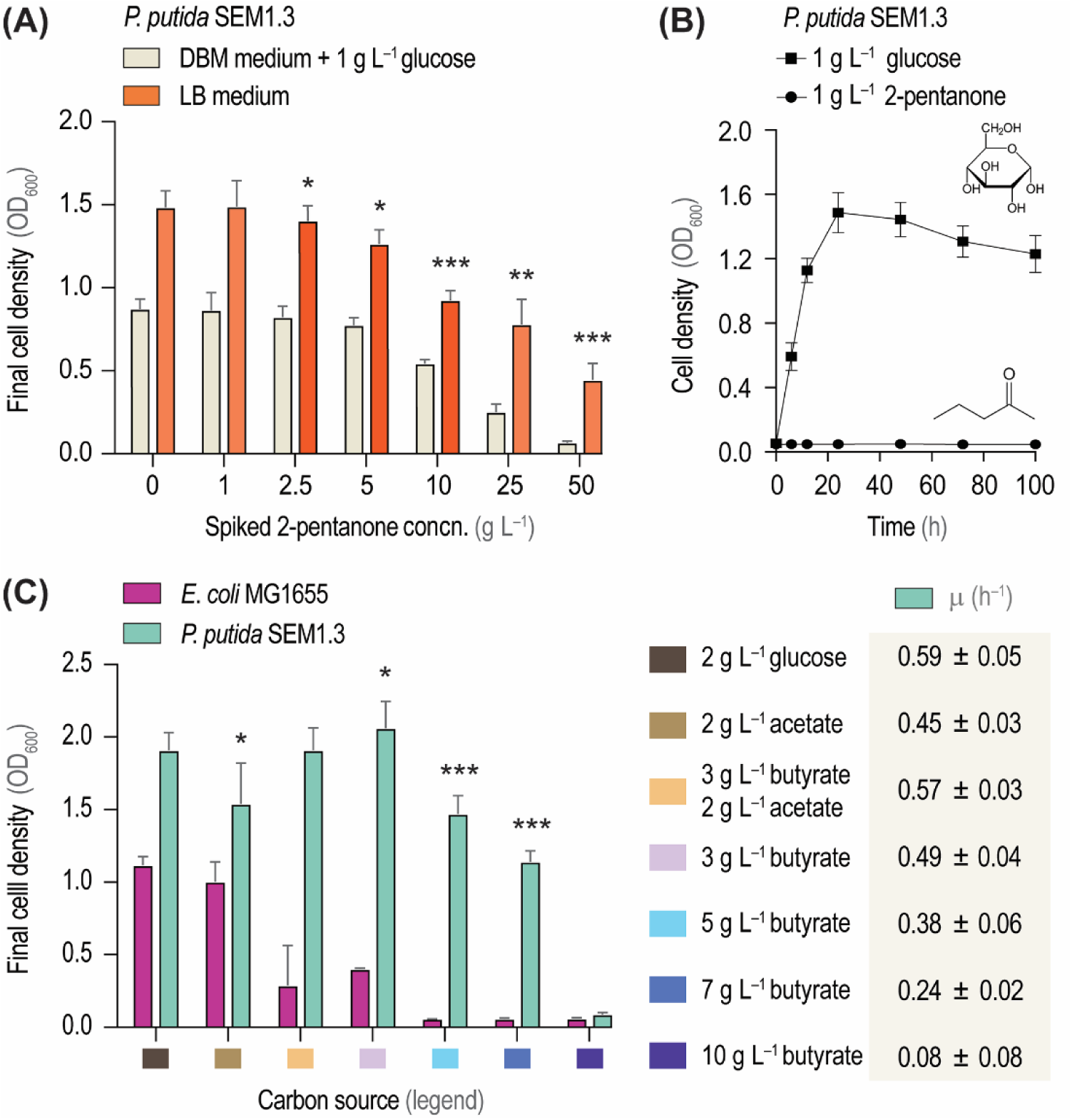
Exploring the tolerance of *P*. *putida* and *E*. *coli* to 2-pentanone and short-chain organic acids. **(A)** Physiological response of *P*. *putida* SEM1.3 to increasing concentrations (concn.) of 2- pentanone, added to either de Bont minimal (DBM) medium or rich LB medium. Cell densities, estimated as the optical density at 600 nm (OD_600_), were measured after 24 h of cultivation. **(B)** Growth profile of *P. putida* SEM1.3 cultivated on DBM medium containing either glucose or 2-pentanone as the sole carbon source. **(C)** Growth of *E. coli* MG1655 and *P. putida* SEM1.3 cultivated in DBM medium with different carbon sources (i.e. glucose, acetate, butyrate or a combination of the two short-chain organic acids) at the concn. indicated. Cell densities, estimated as the OD_600_, were measured after 24 h of cultivation; specific growth rates (μ) were calculated during exponential growth. In all cases, mean values ± standard deviations were derived from independent biological triplicates. Significance levels of final cell density values when compared to control conditions (i.e. 0 g L^−1^ 2-pentanone for panel **A**, 2 g L^−1^ of glucose for panel **B**) are indicated as follows: ∗ *P*-value < 0.05, ** *P*-value < 0.01 and ∗∗∗ *P*- value < 0.001.

**Table 1.**
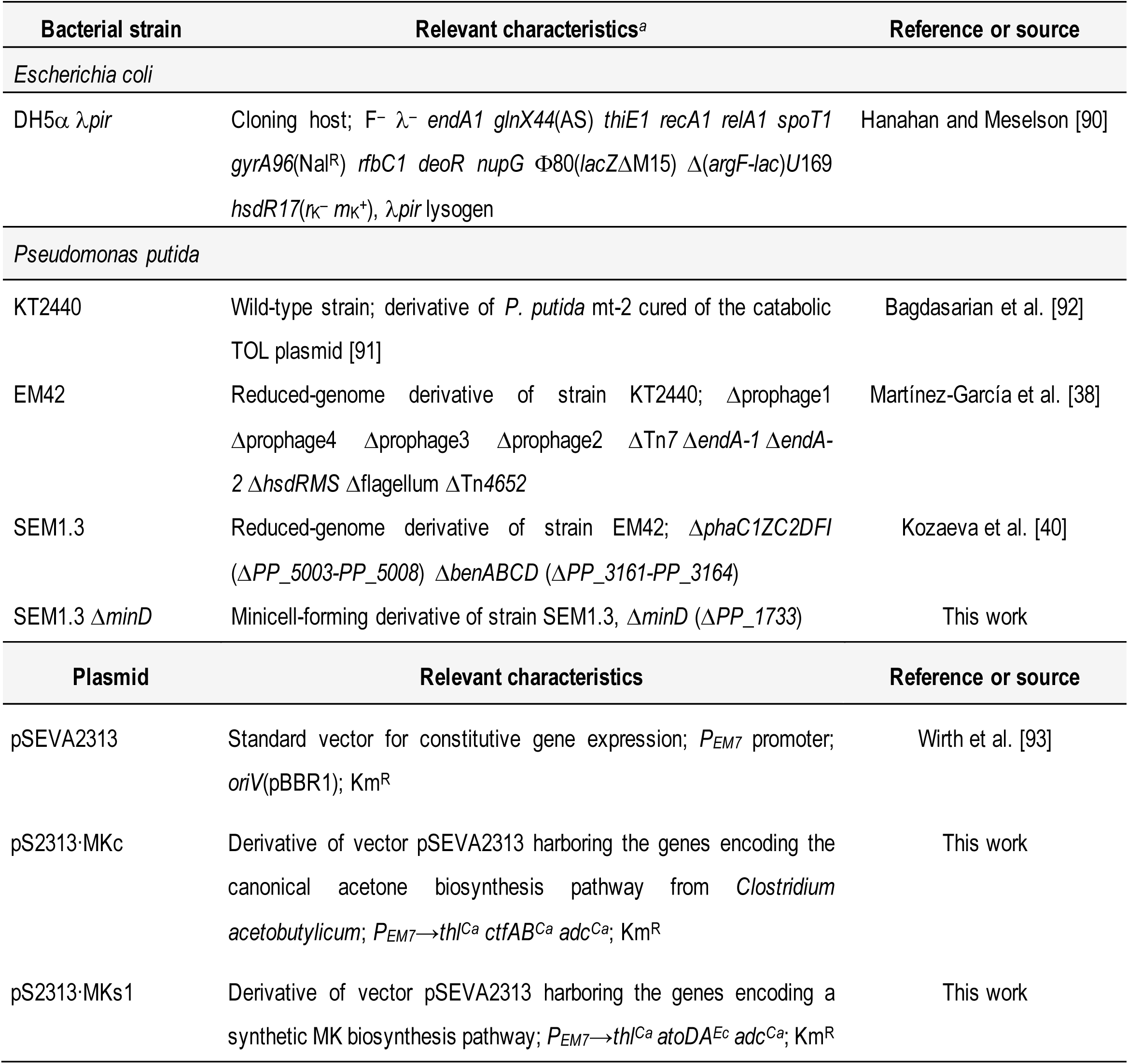
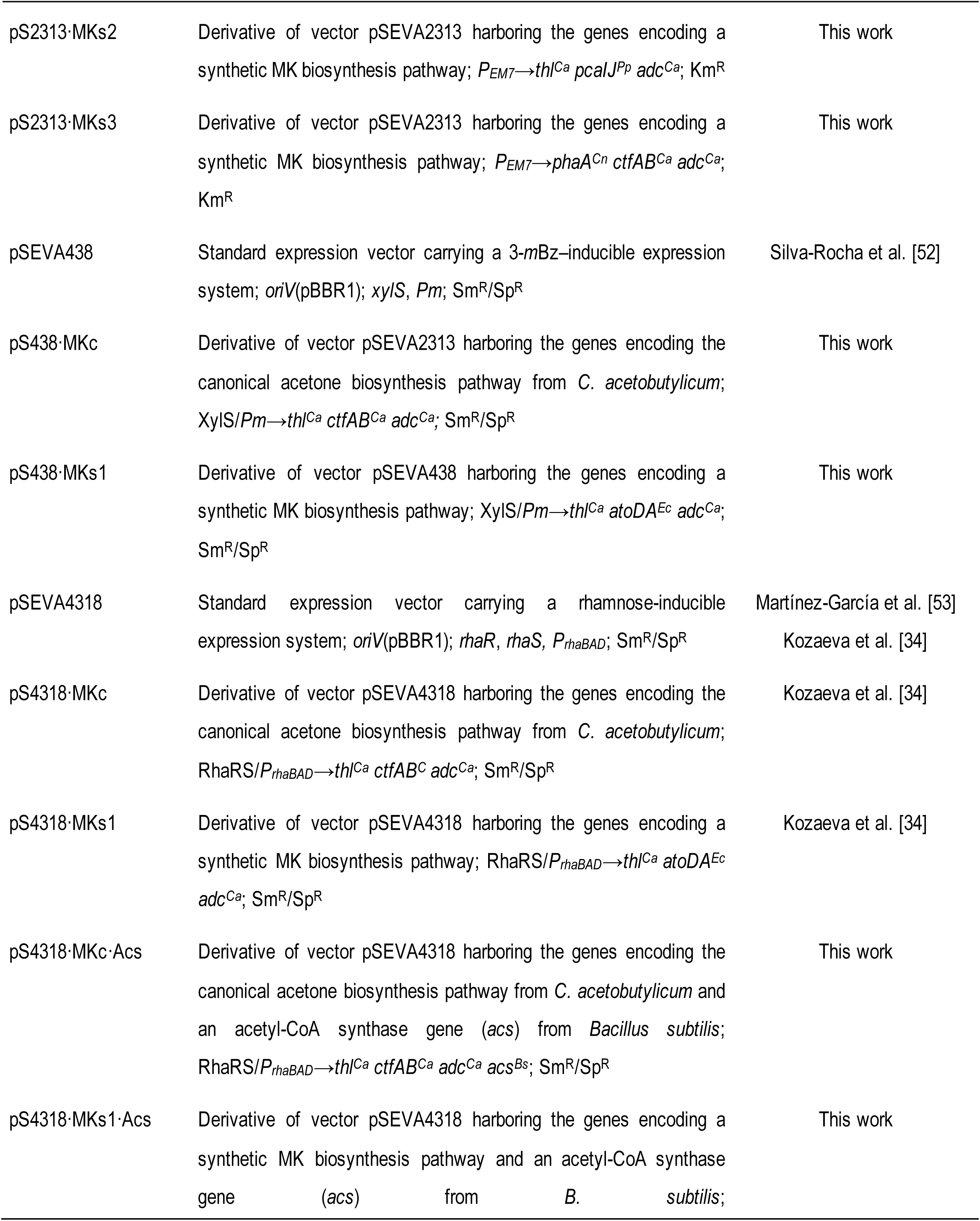
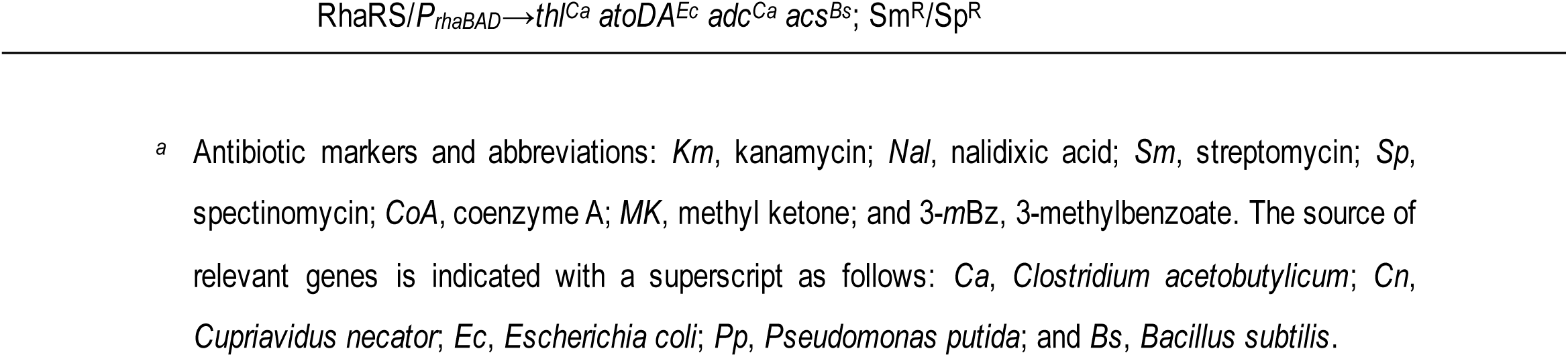
Bacterial strains and plasmids used in this study.

Another attractive metabolic feature of *P. putida*, i.e. efficient assimilation of organic acids as sole carbon source [43] could be exploited for short-chain ketone biosynthesis. To examine this possibility, we evaluated butyrate as a substrate for 2-pentanone production. Butyrate, can be processed by *Pseudomonas* by the canonical β-oxidation pathway to yield acetyl-CoA units [44]. though this carboxylate has not been actively investigated as a feedstock for bacterial fermentations, butyrate is a promising building-block that can be obtained from renewable resources [45] and it has been selected as the substrate of choice for establishing high-cell-density *P. putida* cultures [46]. While these properties position butyrate as an interesting biorefinery substrate, osmotic shock and acid stress may result in toxicity issues even at relatively low concentrations [47]. To test this scenario, the tolerance of *P. putida* SEM1.3 and *E. coli* MG1655 (adopted as a model Gram-negative bacterium extensively used as a host in diverse bioprocesses) [48] to increasing butyrate concentrations was evaluated and compared with widely-used sugar and organic acid substrates (**Fig. 1C**). Glucose or acetate were selected as the primary carbon feedstocks as representative examples of a glycolytic and a gluconeogenic substrate, respectively. *P. putida* SEM1.3 tolerated up to 7 g L^−1^ butyrate with a reduction of ca. 45% in the final OD_600_ values, while the growth of *E. coli* MG1655 was almost abolished at any butyrate concentration above 3 g L^−1^. Neither bacterial species could grow on DBM medium containing 10 g L^−1^ butyrate, which marks the practical upper concentration that can be used in production experiments. Interestingly, the cultures of *P. putida* SEM1.3 reached similar final OD_600_ values when butyrate was either used as the sole carbon substrate or added at 3 g L^−1^ in the presence of glucose or acetate. This observation indicates that butyrate is not a preferred carbon source in the presence of a co-substrate [49]. Moreover, the specific growth rate (μ) was not substantially affected by the addition of butyrate at concentrations < 7 g L^−1^ (**Fig. 1C**). Building on these results, we adopted reduced-genome *P. putida* SEM1.3 as the host to explore the butyrate-dependent biosynthesis of 2- pentanone as explained in the next section.

### Implementing and optimizing pathways for 2-pentanone biosynthesis from organic acids

The canonical short-chain ketone biosynthesis route, key to the widely-known fermentation process to produce acetone by *Clostridium acetobutylicum* ATCC 824 (the *Weizmann organism*) [50] was adopted in this study to explore MK biosynthesis by engineered *P*. *putida*. We hypothesized that this pathway could be coupled with CoA-dependent ketoacid chain elongation [51] thereby yielding butyryl-CoA (C_4_) from acetyl-CoA (C_2_) extender units. In this way, the core acetone biosynthesis pathway of *C. acetobutylicum* can be extended to accommodate the production of a variety of short-chain ketones. As an example, 2-pentanone (C_5_) biosynthesis from butyrate involves the condensation of acetyl-CoA and butyryl-CoA (forming 3-ketohexanoyl-CoA) followed by ester hydrolysis and decarboxylation to yield the final product (**Fig. 2A**). The sequence starts with the condensation of acetyl-CoA and butyryl-CoA mediated by a thiolase (Thl; acetyl-CoA acetyltransferase, EC 2.3.1.9) to generate 3-ketohexanoyl- CoA. Next, CtfAB, an acetoacetyl-CoA:acetate/butyrate CoA transferase (EC 2.8.3.9) relocates a CoA moiety from 3-ketohexanoyl-CoA to acetate, forming 3-ketohexanoate. Finally, this molecule is reduced to 2-pentanone by an acetoacetate decarboxylase (Adc; EC 4.1.1.4), releasing CO_2_ (**Fig. 2A**). Importantly, the theoretical ketone yield from organic acids through the core pathway in **Fig. 2A** is *Y*_P/S_ = 50% mol mol^−1^, which underscores the potential of *P. putida* as a suitable *chassis* for short-chain ketone biosynthesis owing to its tolerance to both substrates and products.

**Fig. 2.**
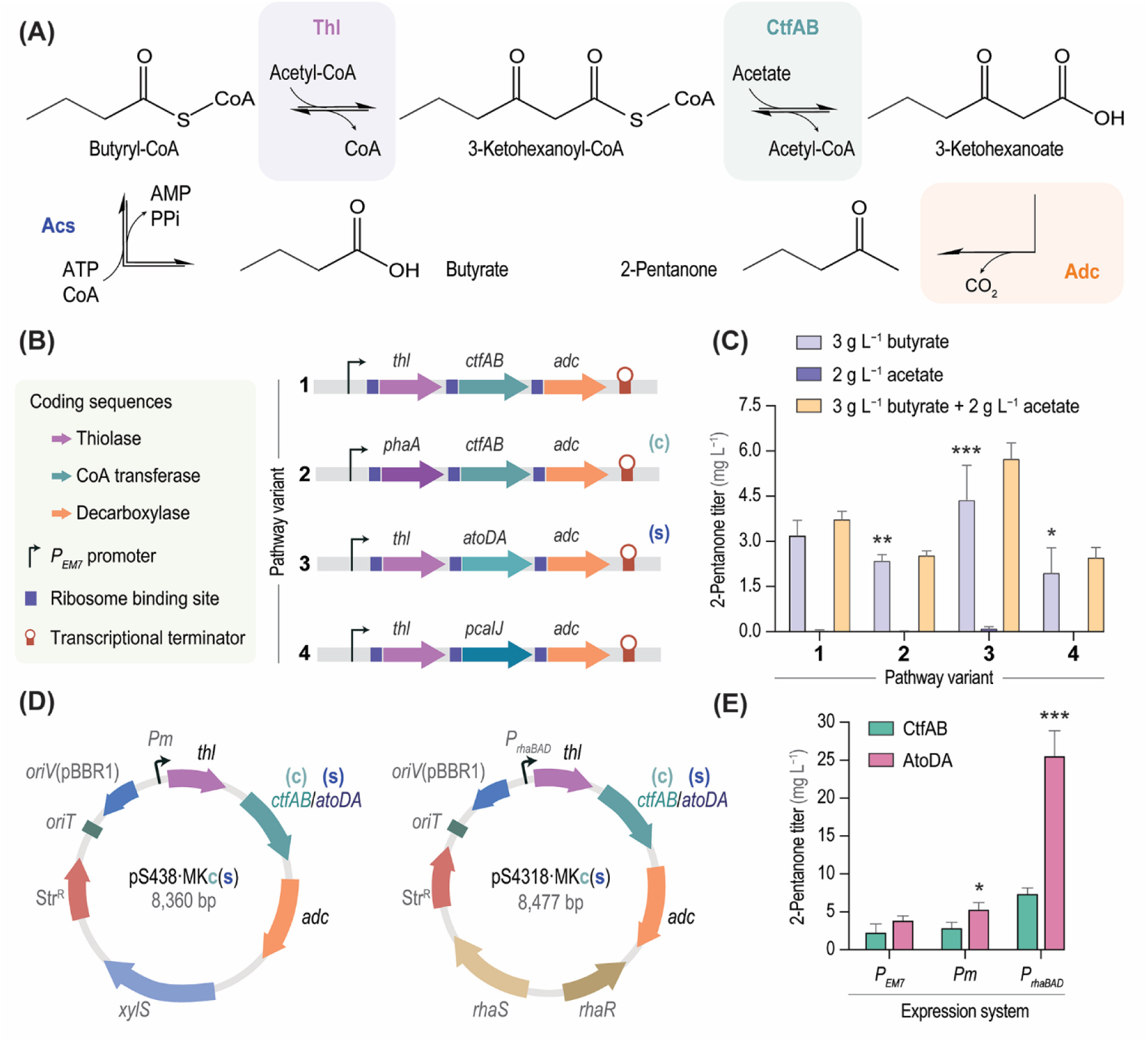
Pathway engineering and optimization for 2-pentanone biosynthesis from short-chain organic acids. **(A)** Biosynthetic pathway for 2-pentanone production from butyrate. Methyl ketone biosynthesis relies on the enzymes of the canonical acetone pathway from *C*. *acetobutylicum* comprising Thl, thiolase (acetyl-CoA acetyltransferase, EC 2.3.1.9), CtfAB (acetoacetyl-CoA:acetate/butyrate CoA transferase, α and β subunits, EC 2.8.3.9) and Adc (acetoacetate decarboxylase, EC 4.1.1.4). Abbreviations: *CoA*, coenzyme A; *PPi*, inorganic pyrophosphate. **(B)** Synthetic operons constructed for 2-pentanone biosynthesis. The elements in this diagram are not drawn to scale. **(C)** Testing 2-pentanone biosynthesis from short-chain organic acids in engineered *P*. *putida*. *P. putida* SEM1.3 was individually transformed with plasmids pS2313·MK[c,s1- s3], carrying the synthetic operons shown in panel **(B)** under control of the constitutive *P_EM7_* promoter (**Table 1**), and incubated for 24 h in DBM medium with the different carbon source combinations indicated; 2-pentanone titers in the culture supernatant were quantified by GC-FID. **(D)** Plasmids constructed for inducible expression of the synthetic operons for 2-pentanone biosynthesis. The synthetic XylS/*Pm* and RhaRS/*P_rhaBAD_* expression systems are induced by addition of 3-methylbenzoate and rhamnose, respectively; both plasmids carry a streptomycin-resistance determinant (Str^R^). **(E)** Exploring strain performance by growing *P. putida* SEM1.3 carrying the selected plasmids in DBM medium containing 3 g L^−1^ butyrate for 24 h. 2-Pentanone titers in the culture supernatant were quantified by GC-FID; methyl ketone titers in strains carrying inducible expression systems are compared to the constitutive expression mediated by the *P_EM7_* promoter. Results shown in panel **(C)** and **(E)** correspond to mean values ± standard deviations from independent biological triplicates. Significance levels of 2-pentanone titers when compared to control conditions (i.e. pathway variant 1, canonical set of enzymes for panel **C;** and constitutive gene expression mediated by the *P_EM7_* promoter for panel **E**) are indicated as follows: ∗ *P*-value < 0.05, ** *P*-value < 0.01 and ∗∗∗ *P*-value < 0.001.

Since all enzymes within the canonical biosynthesis route stem from a Gram-positive bacterium, we decided to expand the biochemical toolset for short-chain ketone production by harnessing activities from species phylogenetically closer to our selected host. To this end, the PhaA thiolase from *Cupriavidus necator* and the AtoDA and PcaIJ CoA transferases from *E. coli* MG1655 and *P. putida* KT2440, respectively, were considered as functional elements for pathway design. The broad-host-range plasmids pS2313·MKc, MKs1, MKs2 and MKs3 (**Table 1**), bearing different combinations of the genes encoding all relevant pathway enzymes under transcriptional control of the constitutive *P_EM7_* promoter, were constructed to this end. A synthetic ribosome binding site [52] (RBS) was incorporated upstream each coding sequence (**Fig. 2B**), and all constructs were inserted into plasmids formatted according to the Standard European Vector Architecture (SEVA) [53]. Note that, according to the plasmid nomenclature adopted in our study, constructs involving canonical pathway components are identified with a *c* letter (e.g. plasmid pS2313·MKc), whereas those bearing synthetic pathway variants are labeled with an *s* letter (e.g. plasmid pS2313·MKs1, where the canonical *ctfAB* genes are replaced by *atoDA* from *E*. *coli*, **Table 1**). The resulting plasmids (pS2313·MKc, MKs1, MKs2 and MKs3) were individually transformed into *P. putida* SEM1.3, and the ability of the engineered strains to produce 2- pentanone was tested in shaken-flask cultures using different carbon substrates. These cultures were grown for 24 h in DBM medium containing either 3 g L^−1^ of butyrate, 2 g L^−1^ of acetate or a combination of 3 g L^−1^ of butyrate and 2 g L^−1^ of acetate as the carbon substrate(s). By the end of the cultivation, 2- pentanone titers and the concentration of any residual carbon source(s) were determined in culture supernatants; the analytic results indicated full consumption of the carbon substrate(s) under all tested conditions. Acetate did not promote MK biosynthesis when used as a sole carbon substrate regardless of the biosynthetic pathway borne by strain SEM1.3 (**Fig. 2C**). Constructs 1 (containing *thl*, *ctfAB* and *adc*) and 3 (spanning *thl*, *atoDA* and *adc*), in contrast, mediated the highest 2-pentanone titers (3.2 mg L^−1^ and 4.5 mg L^−1^, respectively) when butyrate was used as the feedstock. Cultures incubated in the presence of both acetate and butyrate grew to higher cell densities than those added with either organic acid, but the 2-pentanone titers were not significantly different—suggesting that, under these conditions, butyrate was largely responsible of promoting ketone formation. The higher 2-pentanone titers attained by *P*. *putida* SEM13 carrying plasmid pS2313·MKs1 (construct 3) indicates that the acetoacetyl-CoA:acetate/butyrate CoA transferase AtoDA, which moves a CoA moiety from 3- ketohexanoyl-CoA to acetate, is the most efficient enzyme variant to support ketone production. This feature is likely connected to the substrate affinity of AtoDA, which is 23-fold higher than that of CtfAB (*Km* for acetate = 53.1 mM and 1,200 mM, respectively) [54].

Considering the tradeoffs between growth and production [55–57] which compete for essential resources at the level of the acetyl-CoA node [58] we evaluated whether regulated expression of the pathway genes could lead to higher ketone titers. Hence, the pathway optimization process continued by assaying the performance of strain SEM1.3 carrying constructs 1 and 3 under the control of different inducible expression systems. A number of expression systems are available for metabolic engineering of *P. putida*; including both native *Pseudomonas* regulatory elements (e.g. XylS/*Pm*, AlkS/*P_alkB_* and NahR/*P_sal_*) [24] and heterologous modules (e.g. AraC/*P_araB_* and RhaRS/*P_rhaBAD_*) [59]. The XylS/*Pm* and RhaRS/*P_rhaBAD_* expression modules, inducible by 3-methylbenzoate (3-*m*Bz) and rhamnose, respectively, were selected owing to their reportedly tight inducer-dependent regulation features [60] and the high levels of gene expression mediated by these systems [59]. Hence, plasmids pS438·MKc and MKs1 (based on the XylS/*Pm* expression system) and pS4318·MKc and MKs1 (containing the RhaRS/*P_rhaBAD_* module) were constructed for regulated expression of the best-performing pathway designs (constructs 1 and 3, **Table 1**); the physical map of these plasmids is displayed in **Fig. 2D**. These four new plasmids were individually introduced in *P. putida* SEM1.3, and the corresponding cultures were grown for 24 h in DBM medium containing 3 g L^−1^ of butyrate as the carbon substrate, assessing the 2-pentanone titers and residual carbon source concentrations by the end of the incubation period (**Fig. 2E**). The two inducers, 3-*m*Bz and rhamnose, were added at the onset of the cultivation (4 h) at 1 mM and 5 mM, respectively. In general, inducible expression of the different production pathways led to enhanced ketone titers when compared to constitutive gene expression. This positive effect was even more evident in the cultures of *P*. *putida* SEM1.3 carrying construct 3 (which includes the AtoDA transferase), with a 2-pentanone titer of 25.3 mg L^−1^ paired to full substrate consumption.

In the final stage of pathway optimization, it was hypothesized that the activation of butyrate into butyryl-CoA might represent a metabolic bottleneck [61]. Previous experiments depended on the activity of the endogenous acetyl-CoA synthases (Acs) of *P*. *putida* (i.e. Acs-I and Acs-II), necessary for generating the CoA-ester derivatives of the short-chain organic acids used as substrates. To investigate the potential of a heterologous acetyl-CoA synthetase in supporting efficient butyryl-CoA formation, the well-characterized and highly efficient Acs from *Bacillus subtilis* [62] was integrated into the pathway designs, resulting in the construction of plasmids pS4318·MK(c,s1)·*acs* (**Table 1**). However, testing of 2-pentanone biosynthesis by *P*. *putida* SEM1.3 harboring these constructs did not yield a significant enhancement in product titers from butyrate (data not shown). Therefore, it was concluded that the activity of the endogenous Acs-I and Acs-II are probably sufficient for sustaining ketone biosynthesis, and the original pathway designs were retained for further experiments.

### Establishing asymmetrical cell division in P. putida to produce minicells

Building on the innate tolerance of *P*. *putida* to toxic chemicals, we hypothesized that minicells could be harnessed for ketone biosynthesis. As indicated above, minicells are nano-sized (100-400 nm in diameter) achromosomal bacteria, formed by abortive cell division in the mother cell poles [19]. Minicells cannot grow or duplicate, but they can support other vital cellular processes, e.g. ATP synthesis, replication and transcription of plasmid DNA and mRNA translation [19]. To the best of our knowledge, the production and engineering of *P*. *putida* minicells have not been reported thus far, and we explored the cell division mechanisms that could mediate abnormal bacterial segregation in this species. The normal process of cell division in *E*. *coli* and other rod-shaped bacteria, triggered by the formation of the *Z*-ring [18] yields two equally-sized daughter cells [63] (**Fig. 3A**). In Gram-negative bacteria, the dynamics of the *Z*-ring assembly and location are regulated by the Min system [64]. MinD binds to the polar membrane to form a polymer, and the MinCD-complex inhibits FtsZ polymerization of by recruiting MinE—interfering with the membrane assembly at an abnormal location [20]. Abnormal cell division ensues if the *Z*-ring is either formed at the cell pole or if the FtsZ division protein is overproduced, leading to minicell formation [65]. The Min system, thoroughly characterized in *E. coli* and *B. subtilis*, is widespread in bacterial species [15] including *Pseudomonas*. In strain KT2440, *minCDE* form a cluster in the *PP_1732-PP_1734* locus, with the gene encoding the septum site-determining protein MinC as the last element in the sequence [39]. Previous experiments in our laboratory indicated that transcriptional interference on *minD* (*PP_1733*, encoding the ATPase component of the Min system) with a CRISPRi system tailored for *Pseudomonas* species [66] leads to the emergence of minicells from strain KT2440 (data not shown). Hence, an in-frame Δ*minD* deletion mutant of *P. putida* SEM1.3 was created by allelic exchange (**Table 1**). The MinD-deficient *P*. *putida* strain was characterized by scanning electron microscopy (SEM), and a simple three-step protocol, based on sequential centrifugation, was developed for enriching the minicell population (**Fig. 3B**). This procedure consistently yielded ca. 0.25 g_CDW_ (cell dry weight) L^−1^ minicells from a 1.5 g_CDW_ L^−1^ bacterial suspension, and the addition of the broad-spectrum β-lactam ceftriaxone at 100 μg mL^−1^ was sufficient for eliminating any parental cell present in the final minicell preparation. At the last stage of the purification protocol, glycerol was added to the minicell suspension at 15% (v/v) and the preparation was kept at –20°C for extended storage.

**Fig. 3.**
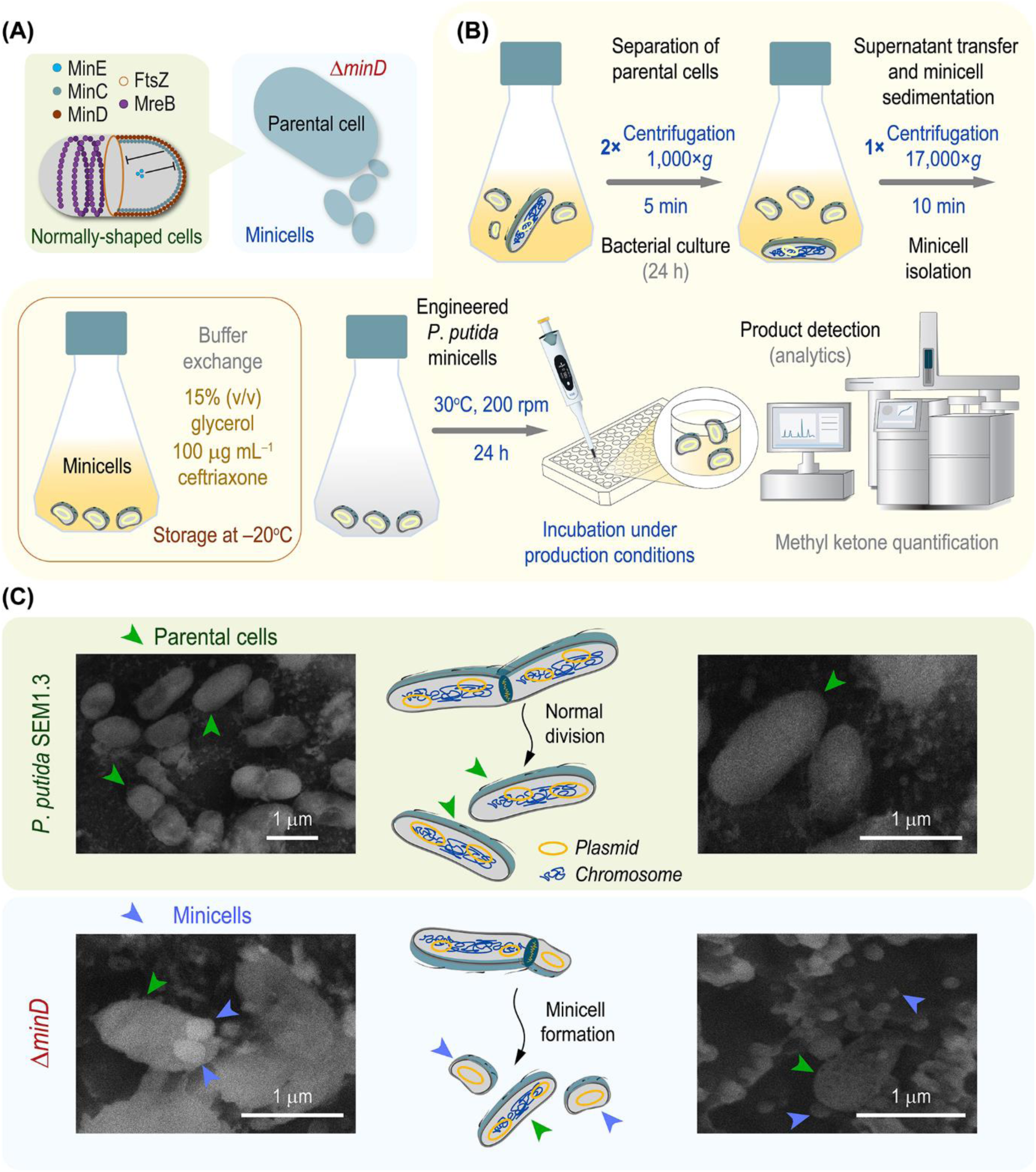
Production and characterization of *P*. *putida* minicells. **(A)** Key elements involved in cell division in Gram-negative bacteria. In normally-shaped cells, the MinC-MinD-MinE system and the cell wall structural, actin-like MreB protein establish a dynamic equilibrium with FtsZ, the division protein, to ensure a proper septation and segregation of daughter cells. Mutations in the Min components, e.g. the *Z*-ring positioning MinD protein, lead to minicell segregation. **(B)** Overview of the protocol implemented for *P*. *putida* minicells production, purification, storage and downstream applications. The resulting minicell suspension can be either used immediately or stored at –20°C (or –70°C) upon buffer exchange. Whenever needed, the minicell preparation is incubated under specific conditions to foster product formation (in this study, acetone and methyl ketones) in microtiter plates, and the metabolites of interest are detected by specific analytical methods. **(C)** Cryo-scanning electron microscopy (SEM) of strains *P*. *putida* SEM1.3 and its Δ*minD* derivative. Cells were grown in 2×YT medium at 30°C with agitation at 150 rpm and harvested after 24 h. Cryo-SEM microscopy was performed on freeze- fractured samples from these cultures, imaged at 15 kV in an X-Max^N^ 150 silicon drift detector (sensor- active area = 150 mm^2^; Oxford Instruments NanoAnalysis) after a Pt-coating treatment. Parental cells and minicells are identified with green and blue arrowheads, respectively; the asymmetrical segregation of daughter cells would lead to the formation of chromosome-free minicells that retain plasmid(s) from the parental *P*. *putida* cell.

To study the minicell-forming phenotype of *P*. *putida* SEM1.3 Δ*minD*, the cells were grown in rich 2×YT medium at 30°C with agitation at 150 rpm and harvested after 24 h. Aliquots of these cultures, and that of the parental strain, were spun down, washed and prepared for cryo-SEM as indicated in *Methods*. Cryo-SEM was performed on freeze-fractured samples derived from these cultures, with imaging at 15 kV in an X-Max^N^ 150 silicon drift detector (**Fig. 3C**). Image analysis identified the segregation of minicells at the poles of *P*. *putida* SEM1.3 Δ*minD* (**Fig. 3C**). The resulting anucleated vesicles were round-shaped, with a mean diameter in the 200-400 nm range, similar to the features reported for *E*. *coli* minicells [67]. In several cases, it was possible to observe minicells budding from normally (rod)- shaped parental bacteria. When the cryo-SEM analysis was conducted again using the minicell- enriched suspension and imaged at a 10,000× magnification, it consistently revealed a uniform population of similarly sized anucleated vesicles (**Fig. 4A**). This result qualitatively validates the effectiveness of the method outlined in **Fig. 3B** for concentrating minicells from cultures of *P*. *putida* SEM1.3 Δ*minD*. Furthermore, *in situ* elemental analysis was conducted for *P*. *putida* SEM1.3 by energy dispersive X-ray spectroscopy (EDS) for carbon (C), nitrogen (N), phosphorus (P) and potassium (K) to assess (whole-cell and minicell) bacterial viability (**Fig. 4B**). SEM-EDS analysis enables semi-quantitative elemental microanalysis by measuring the generation of characteristic X-rays from each chemical element present in the biological specimen, which is particularly relevant when targeting small-sized cells [68]. This analytical technique revealed a clear enrichment of C, N and P in the target area over the background (**Fig. 4B**), whereas the K signal was evenly distributed in the sample, with no clear distinction between the bacterial cells and their surroundings. Hence, the pattern of C signal could be adopted to assess the vitality of bacterial minicells. After validating the reliability of the minicell preparation protocol, the next section focuses on exploring the potential of *P*. *putida* minicells for short-chain ketone bioproduction.

**Fig. 4.**
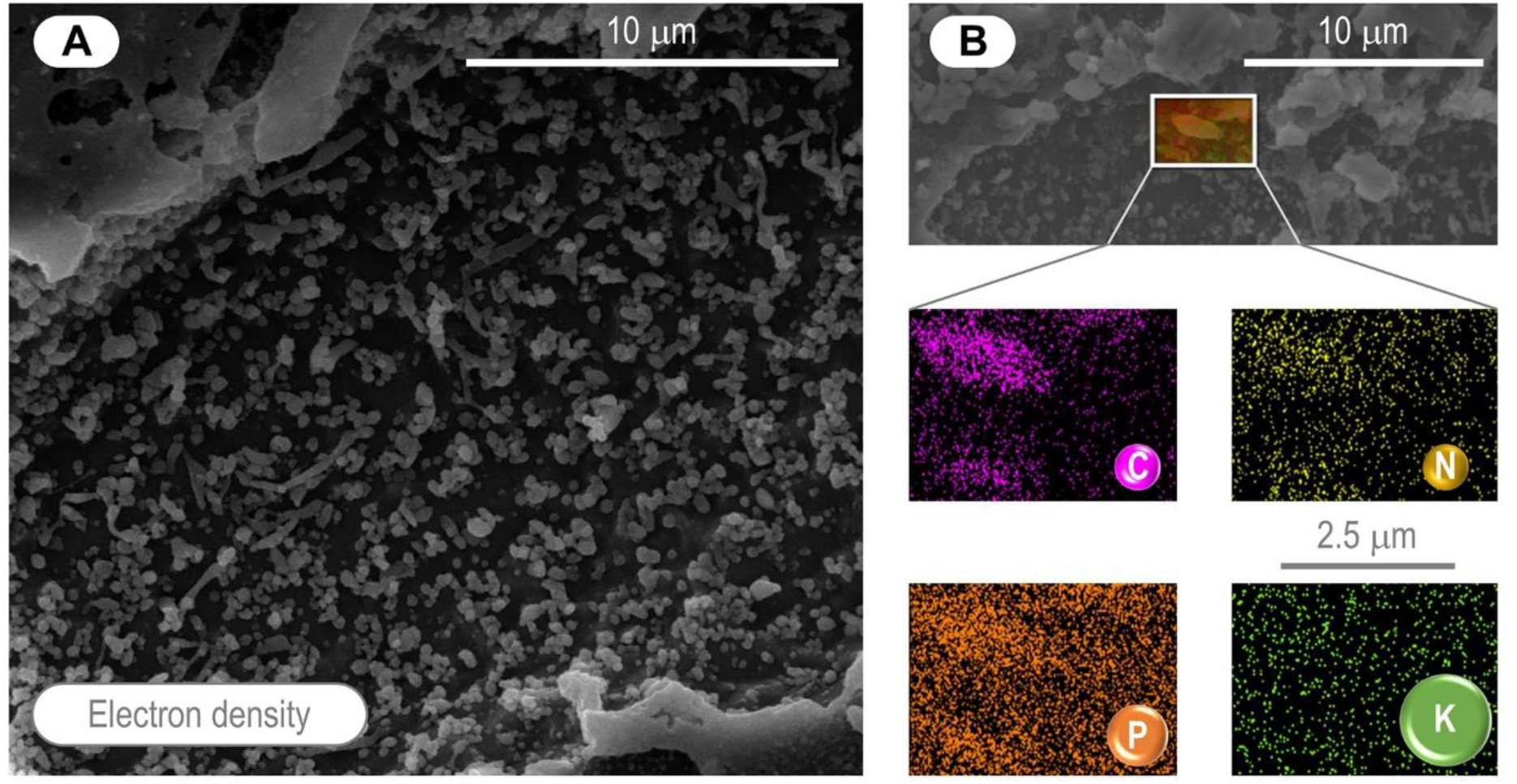
Cryo-scanning electron microscopy (SEM) of *P*. *putida* SEM1.3 and its Δ*minD* (minicell forming) derivative. **(A)** Cryo-SEM frame image-capture of *P*. *putida* minicells. The freeze-fractured, frozen-hydrated sample contains minicells recovered after the last step of purification from cultures of *P*. *putida* SEM1.3 Δ*minD* as indicated in Fig. 3B. The samples were analyzed in an X-Max^N^ 150 silicon drift detector (sensor-active area = 150 mm^2^; Oxford Instruments NanoAnalysis) after a Pt-coating treatment, with magnification = 10,000×. **(B)** *In situ* elemental analysis in a single *P*. *putida* SEM1.3 cell by energy dispersive X-ray spectroscopy (EDS) for carbon (C), nitrogen (N), phosphorus (P) and potassium (K). The SEM-EDS analysis was carried out using two different excitation energies, 5 and 10 kV, to obtain the entire elemental information; the lower excitation energy enabled the detection of N in individual samples. The data for the density maps was collected for 5 min in a single frame image to reduce damage to the sample.

### P. putida minicells mediate bioconversion of organic acids into short-chain ketones

The core pathway designed and optimized for 2-pentanone biosynthesis by engineered *P*. *putida* (borne by plasmid pS4318·MKs1, **Fig. 2**) was transformed into *P*. *putida* SEM1.3 Δ*minD*. We reasoned that the asymmetrical, abortive division in this mutant would lead to the formation of minicells carrying plasmids, but no chromosomal DNA (**Fig. 3C**). Furthermore, we exploited the versatility of the short-chain MK biosynthesis pathway for testing three different products. The same enzymatic cascade, composed by Thl and Adc from *C. acetobutylicum* together with AtoDA from *E. coli*, could be leveraged for the synthesis of various (methyl) ketones by using different, readily available organic acids as feedstocks (e.g. acetate, propanoate and butyrate; **Fig. 5A-C**). Sustainable methods for generating short-chain organic acids are increasingly attracting attention in biotechnology—this trend is exemplified by recent advances in C_1_ assimilation processes for synthesizing building-blocks (e.g. acetate) [69,70]. Acetate, in turn, has been exploited as a feedstock to drive efficient acetone production [71] underscoring the potential of such C_2_ carboxylate as a sustainable carbon source for biorefinery. Hence, by adapting the core biosynthesis pathway (**Fig. 2**), acetone, 2-butanone and 2- pentanone could be produced by feeding acetate (C_2_, **Fig. 5A**), propanoate (C_3_, **Fig. 5B**) and butyrate (C_4_, **Fig. 5C**), respectively, with a theoretical yield *Y*_P/S_ = 50% mol mol^−1^ in all cases.

**Fig. 5.**
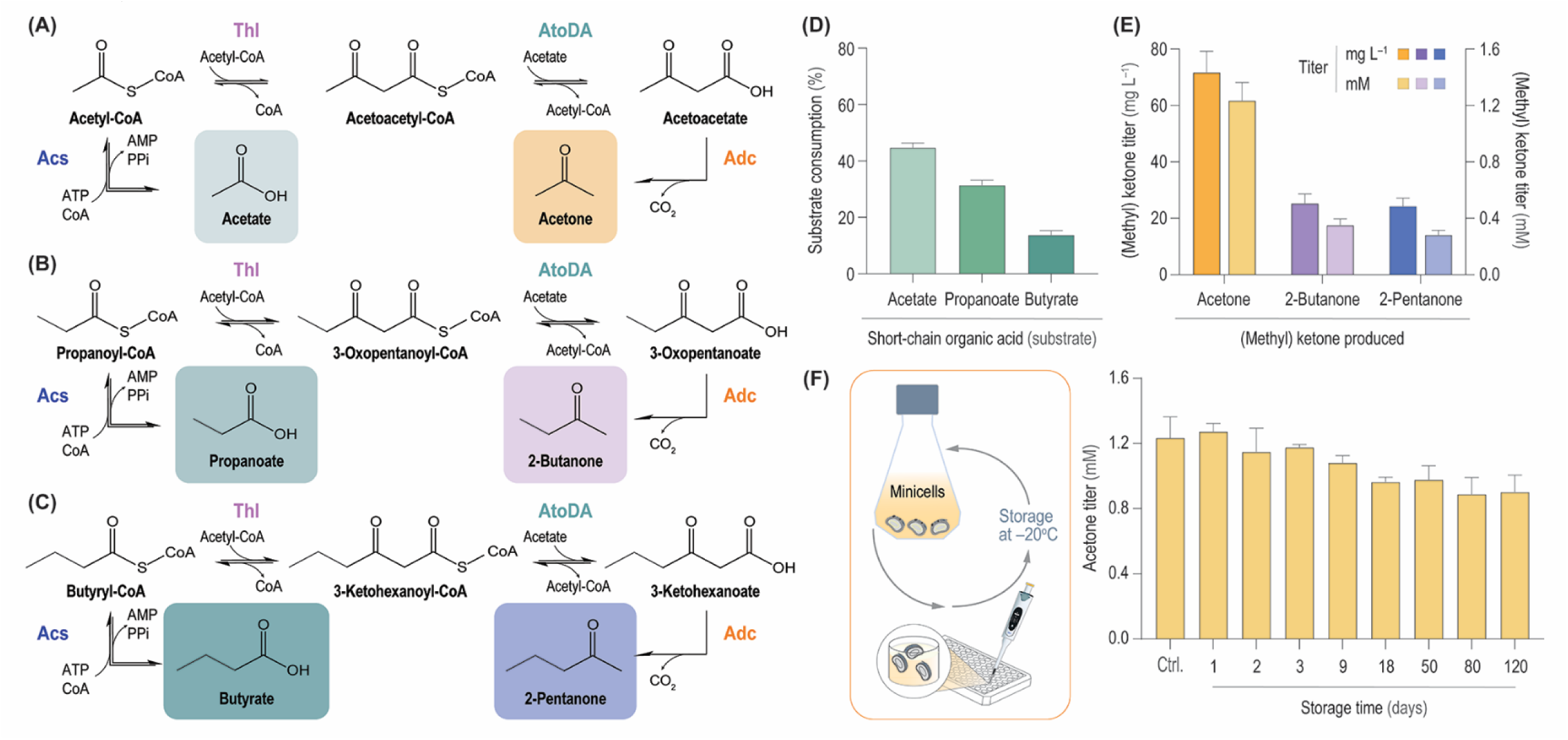
Short-chain (methyl) ketone biosynthesis by *P*. *putida* minicells. **(A-C)** Synthetic pathways tested for the bioconversion of different organic acids into short-chain (methyl) ketones by engineered minicells. The selected pathways mediate the production of acetone from acetate (C _2_) **(A)**, 2-butanone from propanoate (C_3_) **(B)** and 2-pentanone from butyrate (C_4_) **(C)**. The theoretical yield for all ketones in the diagram is *Y_P/S_* = 50% mol mol^−1^; abbreviations: *CoA*, coenzyme A; *PPi*, inorganic pyrophosphate. **(D-E)** Bioconversion experiments using acetate, propanoate or butyrate as the substrate for short-chain ketone production by engineered minicells. In all cases, *P*. *putida* minicells were incubated in the presence of the selected organic acid (50 mM) and both substrate consumption **(D)** and acetone, 2-butanone and 2-pentanone titers **(E)** were determined after 24 h. **(F)** Stability of production phenotypes in *P*. *putida* minicells after prolonged storage. Engineered minicells, prepared as indicated in Fig. 3B, were stored at –20°C (up to 4 months) and periodically tested for their capacity to mediate the bioconversion of acetate into acetone (as indicated in panel **A**). Acetate was used as the substrate at 50 mM, and the acetone titers are compared to a control (Ctrl.) experiment carried out with freshly prepared *P*. *putida* minicells. Results shown in panels **(D-F)** correspond to mean values ± standard deviations from independent biological triplicates.

Minicells derived from *P*. *putida* SEM1.3 Δ*minD* transformed with plasmid pS4318·MKs1 were prepared and purified by following the protocol illustrated in **Fig. 3B**. Bioconversion experiments were established by incubating the same amount of minicells with the three different organic acids under study (i.e. acetate, propanoate and butyrate) at 50 mM. Upon an incubation of 24 h at 30°C with gentle agitation, both substrate consumption and ketone production were assessed in the supernatants. Under these conditions, between 18% and 43% of the carboxylates was consumed by the *P*. *putida* minicells, with a positive correlation between chain length and substrate consumption (**Fig. 4D**). The profile of short-chain MK production by minicells followed the same trend as carboxylate utilization (**Fig. 4E**). Acetone titers reached 72 mg L^−1^ (representing an *Y*_P/S_ value of ca. 20% of the theoretical maximum), whereas the concentration of 2-butanone and 2-pentanone under the same incubation conditions was 25 mg L^−1^ and 22 mg L^−1^, respectively (**Fig. 4E**). Remarkably, the 2-pentanone titers in these experiments were similar to those attained by whole-cell biocatalysis (**Fig. 2E**). These results indicate that the minicells keep the biosynthetic capabilities of the nucleated, parental cells—and the vesicles could produce all three target compounds even when no further engineering steps (e.g. for efficient precursor channeling) has been implemented.

A relatively unexplored advantage of minicells for bioproduction would be catalytic stability, considering that there is neither resource investment nor energy expenditure to support cell division. To probe the stability of *P*. *putida* SEM1.3 Δ*minD* minicells after enrichment and purification (**Fig. 3B**), aliquots of the minicell suspension were stored at –20°C for an extended period. These individual minicell aliquots were periodically retrieved from the freezer and used in acetone bioconversion experiments (24 h) as explained above. The capability of these minicells to convert acetate into the short-chain ketone was analyzed either immediately after minicell purification (serving as the control condition) or after 1, 2, 3, 9, 18, 50, 80 and 120 days of storage (**Fig. 5F**). Acetone titers exhibited little variability across all samples, with >77% of the acetone concentration retained in bioconversion experiments carried out with 4-month-old minicell samples. Moreover, the catalytic activity was kept in minicells stored at –70°C (data not shown), offering further flexibility in the storage conditions. Taken together, these results expose the catalytic stability of *P. putida*-derived minicells, demonstrating their potential to produce toxic chemicals even after prolonged storage.

## CONCLUSION

The biotechnological production of MKs and related molecules is a sustainable alternative to oil-derived chemical production. Here, genome-reduced *P. putida* strains were adapted and optimized for short-chain MK production owing to the innate solvent tolerance and enhanced utilization of organic acids as a carbon source. Examining product and substrate toxicity in both *E. coli* and *P. putida* highlighted the value of *Pseudomonas* as a selected host for MK production. Whole-cell MK production from butyrate as the main carbon source was explored by implementing several pathway designs and synthetic expression systems. This systematic analysis suggested an optimal combination of Thl*^Ca^*, AtoDA*^Ec^* and Adc*^Ca^* functional elements, where the cognate genes are heterologously expressed under the transcriptional control of the rhamnose-inducible RhaRS/*P_rhaBAD_* system. This bioproduction setup is amenable to further optimization steps towards increasing product titers and yields, e.g. fed-batch cultivation [30] or co-feeding strategies [72].

Furthermore, engineered chromosome-free *P. putida* minicells were adopted to explore alternative approaches for short-chain MK bioproduction. *E*. *coli* minicells have been recently exploited for sugar-dependent production of C_6_–C_10_ alcohols and esters [73] and isobutanol and lycopene [74]—leading to titers that often exceeded those attained by whole-cell biocatalysis. In this study, all three MKs of interest (C_3_-C_5_) could be produced by the engineered *P*. *putida* minicells through bioconversion of C*_n_*_–1_ organic acids. Furthermore, the minicell preparation, which can be easily obtained through a 3-step purification protocol, remained catalytically stable even after several weeks of storage in the freezer—a feature that might be further extended up to several months or probably years. Hence, these results not only underscore an alternative bioproduction strategy for chemical commodities (including new-to-nature bioproducts [75]), but also suggest applications for the delivery of chemical cargoes in medical setups. Bacterial minicells can be programmed to target cancer cells [76] and some short-chain ketones have been shown to impair carcinogenesis processes [5] which could be leveraged in medical setups for designing advanced therapies.

## METHODS

### Bacterial strains, plasmids and culture conditions

The bacterial strains and plasmids used in this study are presented in **Table 1**, whereas DNA oligonucleotides and gene fragments are listed in **Table S1** and **S2** in the **Supporting Information**, respectively. *E*. *coli* cultures were incubated at 37°C, *P. putida* cultures were grown at 30°C. Lysogeny broth (LB) medium [77] (10 g L^−1^ tryptone, 5 g L^−1^ yeast extract and 10 g L^−1^ NaCl; solid culture media additionally contained 15 g L^−1^ agar) was employed for cloning procedures and genome engineering manipulations. Rich 2×YT medium (16 g L^−1^ tryptone, 10 g L^−1^ yeast extract and 5 g L^−1^ NaCl) was used to produce *P*. *putida* minicells. In other physiology experiments and unless otherwise specified, de Bont minimal (DBM) medium [78] was used; the culture medium composition is detailed in **Table S3** in the **Supporting Information**. Growth kinetics and related parameters during physiological characterization of engineered strains were derived from measurements of the optical density at 600 nm (OD_600_) with light path correction in a Synergy^TM^ MX microtiter plate reader (BioTek Instruments Inc., Winooski, VT, USA). For whole-cell 2-pentanone production by engineered *P*. *putida* SEM1.3, cultures were initiated from a saturated inoculum, previously grown for 18 h in DBM medium with the same carbon substrate(s) and additives to be used in the main experiment, at a starting OD_600_ of 0.05. Cells were incubated for 24 h at 30°C with constant rotary agitation at 200 rpm (MaxQ™ 8000 shaker incubator; Thermo Fisher Scientific Co., Waltham, MA, USA) in 10 mL of DBM medium contained in a 50-mL Erlenmeyer flask with the corresponding antibiotics and chemical inducers needed for induction of the pathway genes, as indicated in the figure legends. Streptomycin (Sm) and kanamycin (Km) were added whenever needed at 100 μg mL^−1^ and 50 μg mL^−1^, respectively. The OD_600_ values in shaken-flask cultures were periodically recorded in a GENESYS^TM^ 20 spectrophotometer (Thermo Fisher Scientific Co.) to estimate bacterial growth; in all cases, OD_600_ data were processed and analyzed with the QurvE software [79]. The cultures were harvested after 24 h and the flasks were kept at 4°C for 1 h, whereupon the bacterial biomass was recovered by centrifugation at 5,000×*g* for 15 min and 4°C. The concentration of substrates and products in the resulting supernatants was analyzed by HPLC or GC-FID as indicated below.

### General cloning procedures and construction of plasmids and engineered strains

Unless stated otherwise, uracil-excision (*USER*) cloning [80,81] was used for plasmid construction; the AMUSER tool [82] was employed for designing the necessary oligonucleotides. Phusion™ *U* high-fidelity DNA polymerase (Thermo Fisher Scientific Co.) was used according to the manufacturer’s specifications in DNA amplifications intended for *USER* cloning. For colony PCR amplifications, the commercial *OneTaq*™ master mix (New England BioLabs, Ipswich, MA, USA) was used according to the supplier’s instructions. Chemically competent *E. coli* DH5α λ*pir* (**Table 1**) was prepared using the commercially available Mix and Go™ transformation kit (Zymo Research International, Irvin, CA, USA) according to the manufacturer’s indications. These cells were used for plasmid maintenance and other routinary cloning procedures according to well-established molecular biology protocols [83–85]. The sequence of all used plasmids and bacterial strains was verified by Mix2Seq^TM^ sequencing (Eurofins Genomics, Ebersberg, Germany). Plasmids were delivered into *P*. *putida* by electroporating around 300 ng of plasmid DNA into 80 μL of a freshly prepared suspension of electrocompetent cells. *P*. *putida* was rendered electrocompetent by washing four times (by centrifugation and resuspension) the biomass from 4 mL of an overnight culture in LB medium with a 300 mM sucrose solution [86]. Electroporations were performed with a Gene Pulser XCell (Bio-Rad Laboratories Inc., Hercules, CA, USA) set to 2.5 kV, 25 μF capacitance and 200 Ω resistance in a 2-mm gap cuvette.

### Preparation and storage of *P*. *putida* minicells

Precultures for minicell production were grown for 18 h in LB medium containing either 50 μg mL^−1^ Km or 100 μg mL^−1^ Sm (for plasmid-carrying cells), used to inoculate 2×YT medium likewise added with Km or Sm as needed at an initial OD_600_ = 0.05. These 10-mL cultures (placed in 50-mL Falcon^TM^ tubes, Thermo Fisher Scientific Co.) were grown at 30°C with rotary agitation of 150 rpm (Innova™ 42 Incubator Shaker; New Brunswick Scientific Co., Edison, NJ, USA). After 24 h of incubation, the cultures were harvested for minicell purification following the procedure indicated in **Fig. 3B**. After purification, the resulting minicell samples in 50 mM phosphate buffer saline (pH = 7.0) were mixed with glycerol at 15% (v/v) and ceftriaxone at 100 μg mL^−1^ (to prevent bacterial contamination and eliminate any parental cell in the final minicell suspension) towards storing the samples at –20°C or –70°C.

### 2-Pentanone toxicity tests

The tolerance of *P. putida* strains to 2-pentanone was assessed in two experimental setups, in either 50-mL Falcon^TM^ tubes or in microtiter plate cultures, with samples taken every 1 h or at 15-min intervals, respectively, over 72 h. Experiments were performed in biological triplicates using either LB medium or DBM medium containing 1% (w/v) glucose as the main carbon source. Microtiter plates were covered with a Breath-Easy^TM^ sealing membrane (Sigma-Aldrich Co., St. Louis, MO, USA) and OD_600_ readings were recorded automatically under continuous shaking in an Elx808^TM^ absorbance microplate reader (BioTek Instruments Inc.).

### Detection and quantification of MKs and organic acids by HPLC and GC-FID

Culture samples (10 mL) were cooled down in an ice bath and transferred into pre-chilled, 50-mL conical tubes. Culture supernatants, obtained as indicated above, were extracted with hexane (200 μL) and the organic phase was separated by centrifugation for 10 min at 4,500×*g* at 4°C. The organic phase samples were analyzed by gas chromatography (GC) with flame-ionization detection (FID) in a Trace^TM^ 1300 gas chromatograph (Thermo Fisher Scientific Co.) [87–89]. The separation of products was carried out using an Agilent HP-INNOWax capillary column; the results were analyzed using the Chromeleon^TM^ chromatography data system software version 7.1.3 (Thermo Fisher Scientific Co.). Quantification of products was performed by generating a standard calibration curve from the integrated area of spiked samples and calculating the corresponding concentration(s) in experimental samples by the integrated area of their respective peaks. For minicell experiments, the concentrations of acetone, 2-butanone and 2-pentanone as well as that of acetate, propanoate and butyrate were measured by HPLC. A Dionex UltiMate 3000 HPLC system equipped with an Aminex^TM^ HPX-87X ion exclusion (300×7.8 mm) column (Bio-Rad Laboratories Inc.) coupled to a Shodex RI-150 refractive index was used in these measurements. The column was maintained at 30°C, the mobile phase consisted of 5 mM H_2_SO_4_ in Milli-Q water at a flow rate of 0.6 mL min^−1^, with a run length of 45 min. HPLC data were processed using the Chromeleon^TM^ chromatography data system software version 7.1.3 (Thermo Fisher Scientific Co.). The acetone concentration was monitored by refractive index detection, and compound concentrations were calculated from the corresponding peak areas using a calibration curve prepared with authentic standards of acetone, 2-butanone, 2-pentanone, sodium acetate, sodium propanoate and sodium butyrate (in all cases, >99% HPLC standards were used; Sigma-Aldrich Co.).

### Synthesis of short-chain MKs in bioconversion experiments using *P*. *putida* minicells

Production of short-chain ketones was assayed in minicells using different carboxylates as substrates. Sodium acetate, sodium propanoate and sodium butyrate were selected to evaluate the production of acetone, 2-butanone and 2-pentanone, respectively. The selected carboxylate was added to 235 μL of the minicell suspension (ca. 0.5 g_CDW_ L^−1^) at a concentration of 50 mM (final volume = 240 μL); the mixtures were incubated at 30°C with rotary agitation at 180 rpm (Innova™ 42 Incubator Shaker) for 24 h. The formation of the corresponding short-chain ketones was analyzed by HPLC and calculated as described above; equivalent assays were performed in parallel to study the consumption of the selected carboxylates.

### Cryo-scanning electron microscopy (cryo-SEM) for minicell visualization and characterization

Cryo-SEM was done to observe morphology of individual cells in bacterial cultures of *P*. *putida* SEM1.3 and its Δ*minD* derivative. Cells were grown in 2×YT medium at 30°C with agitation at 150 rpm and harvested after 24 h, the OD_600_ of every sample was normalized to 1, and purification of minicells was done as indicated in **Fig. 3B**. The protocol consisted of the following steps: (i) the parental *P*. *putida* cells were sedimented by centrifugation at low speed (5 min at 1,000×*g* and 4°C, repeated 2 times), (ii) the supernatant was carefully transferred into a new Falcon^TM^ tube and centrifuged at 17,000×*g* to recover *P*. *putida* minicells and (iii) the resulting minicell pellet was resuspended in 5 mL of the buffer of choice (typically, 50 mM phosphate buffered saline, pH = 7.0). For quality control and phenotype validation, microscopy analysis was performed using cryo-scanning electron microscopy (SEM) of a freeze-fractured sample, coated with gold, and imaged at 15 kV in an X-Max^N^ 150 silicon drift detector (sensor-active area = 150 mm^2^; Oxford Instruments NanoAnalysis, Abingdon, United Kingdom). The bead was mounted for cryo-SEM experiments on a sample holder attached to a transfer rod, rapidly frozen by plunging into slushed liquid nitrogen (N_2_) at −210°C and transferred to the preparation chamber stage at −180°C (PP2000 CryoTransfer^TM^ System; Quorum Technologies, Laughton, East Sussex, United Kingdom). Frozen samples were cleaved with a cold knife (facilitating an exposed surface in the fractured sample), sublimated at −80°C for 15 min and coated with Pt at a current of 4.5 mA for 30 s. The samples were then transferred under vacuum to the SEM stage in a field-emission scanning electron microscope (Quanta^TM^ 200 FEG; FEI Co., Hillsboro, OR, USA) and imaged at 10 kV using an Everhart–Thornley detector. The pore size distribution of beads was analyzed using the ImageJ software (version 1.50b; National Institutes of Health, Bethesda, MD, USA).

### Data and statistical analysis

All the experiments reported were independently repeated at least in three independent biological replicates (as indicated in the corresponding figure legend), and the mean value of the corresponding parameter ± standard deviation is presented. When relevant, the level of significance of differences when comparing results was evaluated by ANOVA (Barlett’s test) with α = 0.001, 0.01 or 0.05, as indicated in the figure legends. Data analysis was performed with MS Excel^TM^ (Microsoft Corp., Redmond, WA, USA) and Prism 8 (GraphPad Software Inc., San Diego, CA, USA) unless differently specified.

## SUPPORTING INFORMATION

**Table S1.** Oligonucleotides used in this work.

**Table S2.** Gene fragments used in this work.

**Table S3**. Composition of de Bont minimal medium.

## ACKNOWLEDGMENTS

We are indebted to the Microscopy Team, especially the characterization specialist Marie Karen Tracy Hong Lin at DTU NanoLab (National Centre for Nano Fabrication and Characterization, Technical University of Denmark), for their help with processing of the samples and visualization by cryo-SEM. E.K. was the recipient of a fellowship from The Novo Nordisk Foundation as part of the Copenhagen Bioscience Ph.D. Programme (Cohort 2017), supported through grant NNF17CC0026768. M.N.D acknowledges the support received from the European Union’s *Horizon2020* Research and innovation programme under the Marie Sklodowska-Curie grant agreement No. 713683 (*COFUNDfellowsDTU*) and from the VELUX Foundation under the Villum Experiment program (grant No. 40979). The financial support from The Novo Nordisk Foundation (grants NNF18OC0034818, NNF20CC0035580 and NNF21OC0067996) and the European Union’s Horizon2020 Research and Innovation Programme under grant agreement No. 814418 (*SinFonia*) to P.I.N. is also gratefully acknowledged. The authors declare that there are no competing interests associated with the contents of this article.

## AUTHOR CONTRIBUTIONS

**E.K.:** Conceptualization, Investigation, Formal analysis, Validation, Supervision, Methodology, Visualization, Writing─original draft; **M.N.D:** Investigation, Formal analysis, Validation, Methodology, Visualization; **K.K.Y.T:** Investigation, Methodology; **P.I.N:** Conceptualization, Resources, Supervision, Project administration, Funding acquisition, Writing─review and editing, Visualization, Data curation.

