## Supporting Information for "Leveraging engineered *Pseudomonas putida* minicells for bioconversion of organic acids into short-chain methyl ketones"

by

Ekaterina Kozaeva†, Manuel Nieto-Domínguez, Kent Kang Yong Tang‡, and Pablo Iván Nikel*

The Novo Nordisk Foundation Center for Biosustainability, Technical University of Denmark, 2800 Kongens Lyngby, Denmark

† *Current address*: Department of Bioengineering, Stanford University, Stanford, CA, USA
‡ *Current address*: Novo Nordisk A/S, Hovedstaden, Denmark

**Table S1.** Oligonucleotides used in this work.

| # | Name | Sequence (5'→3') | Application |
|---|---|---|---|
| 1 | U-SEVA43-F | ATCGCCCUAGGCCGCGGCCGCGCGAATTCGA | USER cloning of elements of the rhamnose-inducible expression system (RhaRS/$P_{rhaBAD}$), to build vector pSEVA4318 and derivatives |
| 2 | U-SEVA43-R | AAAGGCAUCAAATAAAACGAAAGGCTCAGTC | |
| 3 | U-*rhaSR*-F | ATGCCTTUAATTAAAGCGGATAACAATTTCAC | |
| 4 | U-*rhaSR*-R | AGGGCGAUCGCCTCAGCGAATTTCATTACGA | |
| 5 | U-SEVA43R-F | AGGCGTCGUGACTGGGAAAACCCTGGCG | USER cloning to assemble a canonical production pathway, comprising *thl, ctfAB* and *adc*; synthetic RBSs for each gene were added in the oligonucleotide sequences |
| 6 | U-SEVA43R-R | ACCTCCUACCGCGGCCTAGGGCGA | |
| 7 | U-*thl*-F | AGGAGGUTAGTTAGAATGAAAGAAG | |
| 8 | U-*thl*-R | TTCAUTTTAATCCCTCCTTTTCTAGC | |
| 9 | U-*ctfAB*-F | AAAAUGAACTCTAAAATAATTAGATT | |
| 10 | U-*ctfAB*-R | ACAGCCAUGGGTCTAAGTTCATTGG | |
| 11 | U-*adc*-F | ATGGCTGUTTAGTTTAGGAAGGTGACT | |
| 12 | U-*adc*-R | ACGACGCCUTACTTAAGATAATCATATATAA | |
| 13 | U-S43R-*adc*-F | ATTTATGAUTTAGGAAGGTGACTTTT | USER cloning to exchange *ctfAB* with *atoDA* genes to build a synthetic production pathway |
| 14 | U-S43R-*thl*-R | AGCACTTUTCTAGCAATATTGCTGT | |
| 15 | U-*atoDA*-for | ATCATAAAUCACCCCGTTGCGTATT | |
| 16 | U-*atoDA*-rev | AGAAAAGUGCTAGAAAAGGAGGGATTA | |
| 17 | U-SEVA23-F | ATCGCCCUAGGCCGCGGCCGCGCGAATCGA | USER cloning to assemble production pathways with constitutive expression |
| 18 | U-SEVA23-R | AAAGGCAUCAAATAAAACGAAAGGCTCAGTC | |
| 19 | U-$P_{EM7}$-F | ATGCCTTUAATTAAAGCGGATAACAATCAC | |
| 20 | U-$P_{EM7}$-R | AGGGCGAUCGCCTCAGCGAATTTCATTACG | |

| | | | |
|---|---|---|---|
| 21 | U-SEVA43-F | ATCGCCCUAGGCCGCGGCCGCGCGAA | USER cloning to assemble production pathways with the XylS/*Pm* expression system |
| 22 | U-SEVA43-R | AAAGGCAUCAAATAAAACGAAAGGCTCAGTC | |
| 23 | U-*xylS*-F | ATGCCTTUAATTAAAGCGGATAACTTCAC | |
| 24 | U-*xylS*-R | AGGGCGAUCGCCTCAGCGAATTTCTACGA | |
| 25 | U-*phaA*-F | ATCGCCCUAGGCCGCGGCCGCGCGAATCGA | USER cloning to exchange *thl* with *phaA* gene |
| 26 | U-*phaA*-R | AAAGGCAUCAAATAAAACGAAAGGCTCA | |

**Table S2.** Gene fragments used in this work.

| Name | Sequence (5'→3') |
|---|---|
| Sequence encoding *thl* gene | **ATG**AAAGAAGTTGTAATAGCTAGTGCAGTAAGAACAGCGATTGGATCTTATGGAAA<br>GTCTCTTAAGGATGTACCAGCAGTAGATTTAGGAGCTACAGCTATAAAGGAAGCAG<br>TTAAAAAAGCAGGAATAAAACCAGAGGATGTTAATGAAGTCATTTTAGGAAATGTT<br>CTTCAAGCAGGTTTAGGACAGAATCCAGCAAGACAGGCATCTTTTAAAGCAGGATT<br>ACCAGTTGAAATTCCAGCTATGACTATTAATAAGGTTTGTGGTTCAGGACTTAGAA<br>CAGTTAGCTTAGCAGCACAAATTATAAAAGCAGGAGATGCTGACGTAATAATAGCA<br>GGTGGTATGGAAAATATGTCTAGAGCTCCTTACTTAGCGAATAACGCTAGATGGGG<br>ATATAGAATGGGAAACGCTAAATTTGTTGATGAAATGATCACTGACGGATTGTGGG<br>ATGCATTTAATGATTACCACATGGGAATAACAGCAGAAAACATAGCTGAGAGATGG<br>AACATTTCAAGAGAAGAACAAGATGAGTTTGCTCTTGCATCACAAAAAAAAGCTGA<br>AGAAGCTATAAAATCAGGTCAATTTAAAGATGAAATAGTTCCTGTAGTAATTAAAG<br>GCAGAAAGGGAGAAACTGTAGTTGATACAGATGAGCACCCTAGATTTGGATCAACT<br>ATAGAAGGACTTGCAAAATTAAAACCTGCCTTCAAAAAAGATGGAACAGTTACAGC<br>TGGTAATGCATCAGGATTAAATGACTGTGCAGCAGTACTTGTAATCATGAGTGCAG<br>AAAAAGCTAAAGAGCTTGGAGTAAAACCACTTGCTAAGATAGTTTCTTATGGTTCA<br>GCAGGAGTTGACCCAGCAATAATGGGATATGGACCTTTCTATGCAACAAAAGCAGC<br>TATTGAAAAAGCAGGTTGGACAGTTGATGAATTAGATTTAATAGAATCAAATGAAG<br>CTTTTGCAGCTCAAAGTTTAGCAGTAGCAAAAGATTTAAAATTTGATATGAATAAA<br>GTAAATGTAAATGGAGGAGCTATTGCCCTTGGTCATCCAATTGGAGCATCAGGTGC<br>AAGAATACTCGTTACTCTTGTACACGCAATGCAAAAAAGAGATGCAAAAAAAGGCT<br>TAGCAACTTTATGTATAGGTGGCGGACAAGGAACAGCAATATTGCTAGAAAAGTGC<br>**TAG** |
| Sequence encoding *ctfAB* genes | **ATG**AACTCTAAAATAATTAGATTTGAAAATTTAAGGTCATTCTTTAAAGATGGGAT<br>GACAATTATGATTGGAGGTTTTTTAAACTGTGGCACTCCAACCAAATTAATTGATT<br>TTTTAGTTAATTTAAATATAAAGAATTTAACGATTATAAGTAATGATACATGTTAT<br>CCTAATACAGGTATTGGTAAGTTAATATCAAATAATCAAGTAAAAAAGCTTATTGC<br>TTCATATATAGGCAGCAACCCAGATACTGGCAAAAAACTTTTTAATAATGAACTTG<br>AAGTAGAGCTCTCTCCCCAAGGAACTCTAGTGGAAAGAATACGTGCAGGCGGATCT<br>GGCTTAGGTGGTGTACTAACTAAAACAGGTTTAGGAACTTTGATTGAAAAAGGAAA<br>GAAAAAAATATCTATAAATGGAACGGAATATTTGTTAGAGCTACCTCTTACAGCCG<br>ATGTAGCATTAATTAAAGGTAGTATTGTAGATGAGGCCGGAAACACCTTCTATAAA<br>GGTACTACTAAAAACTTTAATCCCTATATGGCAATGGCAGCTAAAACCGTAATAGT<br>TGAAGCTGAAAATTTAGTTAGCTGTGAAAAACTAGAAAAGGAAAAAGCAATGACCC<br>CCGGAGTTCTTATAAATTATATAGTAAAGGAGCCTGCA**TAA**A**ATG**ATTAATGATAA<br>AAACCTAGCGAAAGAAATAATAGCCAAAAGAGTTGCAAGAGAATTAAAAAATGGTC<br>AACTTGTAAACTTAGGTGTAGGTCTTCCTACCATGGTTGCAGATTATATACCAAAA<br>AATTTCAAAATTACTTTCCAATCAGAAAACGGAATAGTTGGAATGGGCGCTAGTCC<br>TAAAATAAATGAGGCAGATAAAGATGTAGTAAATGCAGGAGGAGACTATACAACAG<br>TACTTCCTGACGGCACATTTTTCGATAGCTCAGTTTCGTTTTCACTAATCCGTGGT<br>GGTCACGTAGATGTTACTGTTTTAGGGGCTCTCCAGGTAGATGAAAAGGGTAATAT<br>AGCCAATTGGATTGTTCCTGGAAAAATGCTCTCTGGTATGGGTGGAGCTATGGATT<br>TAGTAAATGGAGCTAAGAAAGTAATAATTGCAATGAGACATACAAATAAAGGTCAA<br>CCTAAAATTTTAAAAAAATGTACACTTCCCCTCACGGCAAAGTCTCAAGCAAATCT<br>AATTGTAACAGAACTTGGAGTAATTGAGGTTATTAATGATGGTTTACTTCTCACTG<br>AAATTAATAAAAACACAACCATTGATGAAATAAGGTCTTTAACTGCTGCAGATTTA<br>CTCATATCCAATGAACTTAGACCCATGGCTGTT**TAG** |
| Sequence encoding *atoDA* genes | **ATG**AAAACAAAATTGATGACATTACAAGACGCCACCGGCTTCTTTCGTGACGGCAT<br>GACCATCATGGTGGGCGGATTTATGGGGATTGGCACTCCATCCCGCCTGGTTGAAG<br>CATTACTGGAATCTGGTGTTCGCGACCTGACATTGATAGCCAATGATACCGCGTTT<br>GTTGATACCGGCATCGGTCCGCTCATCGTCAATGGTCGAGTCCGCAAAGTGATTGC<br>TTCACATATCGGCACCAACCCGGAAACAGGTCGGCGCATGATATCTGGTGAGATGG<br>ACGTCGTTCTGGTGCCGCAAGGTACGCTAATCGAGCAAATTCGCTGTGGTGGAGCT |

| | |
|---|---|
| | GGACTTGGTGGTTTTCTCACCCCAACGGGTGTCGGCACCGTCGTAGAGGAAGGCAA<br>ACAGACACTGACACTCGACGGTAAAACCTGGCTGCTCGAACGCCCACTGCGCGCCG<br>ACCTGGCGCTAATTCGCGCTCATCGTTGCGACACACTTGGCAACCTGACCTATCAA<br>CTTAGCGCCCGCAACTTTAACCCCCTGATAGCCCTTGCGGCTGATATCACGCTGGT<br>AGAGCCAGATGAACTGGTCGAAACCGGCGAGCTGCAACCTGACCATATTGTCACCC<br>CTGGTGCCGTTATCGACCACATCATCGTTTCACAGGAGAGCAAA**TAATG**GATGCGA<br>AACAACGTATTGCGCGCCGTGTGGCGCAAGAGCTTCGTGATGGTGACATCGTTAAC<br>TTAGGGATCGGTTTACCCACAATGGTCGCCAATTATTTACCGGAGGGTATTCATAT<br>CACTCTGCAATCGGAAAACGGCTTCCTCGGTTTAGGCCCGGTCACGACAGCGCATC<br>CAGATCTGGTGAACGCTGGCGGGCAACCGTGCGGTGTTTTACCCGGTGCAGCCATG<br>TTTGATAGCGCCATGTCATTTGCGCTAATCCGTGGCGGTCATATTGATGCCTGCGT<br>GCTCGGCGGTTTGCAAGTAGACGAAGAAGCAAACCTCGCGAACTGGGTAGTGCCTG<br>GGAAAATGGTGCCCGGTATGGGTGGCGCGATGGATCTGGTGACCGGGTCGCGCAAA<br>GTGATCATCGCCATGGAACATTGCGCCAAAGATGGTTCAGCAAAAATTTTGCGCCG<br>CTGCACCATGCCACTCACTGCGCAACATGCGGTGCATATGCTGGTTACTGAACTGG<br>CTGTCTTTCGTTTTATTGACGGCAAAATGTGGCTCACCGAAATTGCCGACGGGTGT<br>GATTTAGCCACCGTGCGTGCCAAAACAGAAGCTCGGTTTGAAGTCGCCGCCGATCT<br>GAATACGCAACGGGGTGATTTA**TGA** |
| Sequence encoding *adc* gene | **ATG**TTAAAGGATGAAGTAATTAAACAAATTAGCACGCCATTAACTTCGCCTGCATT<br>TCCTAGAGGACCCTATAAATTTCATAATCGTGAGTATTTTAACATTGTATATCGTA<br>CAGATATGGATGCACTTCGTAAAGTTGTGCCAGAGCCTTTAGAAATTGATGAGCCC<br>TTAGTCAGGTTTGAAATTATGGCAATGCATGATACGAGTGGACTTGGTTGTTATAC<br>AGAAAGCGGACAGGCTATTCCCGTAAGCTTTAATGGAGTTAAGGGAGATTATCTTC<br>ATATGATGTATTTAGATAATGAGCCTGCAATTGCAGTAGGAAGGGAATTAAGTGCA<br>TATCCTAAAAAGCTCGGGTATCCAAAGCTTTTTGTGGATTCAGATACTTTAGTAGG<br>AACTTTAGACTATGGAAAACTTAGAGTTGCGACAGCTACAATGGGGTACAAACATA<br>AAGCCTTAGATGCTAATGAAGCAAAGGATCAAATTTGTCGCCCTAATTATATGTTG<br>AAAATAATACCCAATTATGATGGAAGCCCTAGAATATGTGAGCTTATAAATGCGAA<br>AATCACAGATGTTACCGTACATGAAGCTTGGACAGGACCAACTCGACTGCAGTTAT<br>TTGATCACGCTATGGCGCCACTTAATGATTTGCCAGTAAAAGAGATTGTTTCTAGC<br>TCTCACATTCTTGCAGATATAATATTGCCTAGAGCTGAAGTTATATATGATTATCT<br>TAAG**TAA** |

**Table S3**. Composition of de Bont minimal medium.

| Component | Final concentration |
|---|---|
| $K_2HPO_4$ | 3.88 g $L^{-1}$ |
| $NaH_2PO_4$ | 1.63 g $L^{-1}$ |
| $(NH_4)_2SO_4$ | 2 g $L^{-1}$ |
| $MgCl_2·6H_2O$ | 0.1 g $L^{-1}$ |
| EDTA | 10 mg $L^{-1}$ |
| $ZnSO_4·7H_2O$ | 2 mg $L^{-1}$ |
| $CaCl_2·2H_2O$ | 1 mg $L^{-1}$ |
| $FeSO_4·7H_2O$ | 5 mg $L^{-1}$ |
| $Na_2MoO_4·2H_2O$ | 0.2 mg $L^{-1}$ |
| $CuSO_4·5H_2O$ | 0.2 mg $L^{-1}$ |
| $CoCl_2·6H_2O$ | 0.4 mg $L^{-1}$ |
| $MnCl_2·2H_2O$ | 1 mg $L^{-1}$ |
